# Development and clinical validation of a targeted RNAseq panel (Fusion-STAMP) for diagnostic and predictive gene fusion detection in solid tumors

**DOI:** 10.1101/870634

**Authors:** Erik Nohr, Christian A. Kunder, Carol Jones, Shirley Sutton, Eula Fung, Hongbo Zhu, Sharon J. Feng, Linda Gojenola, Carlos D. Bustamante, James L. Zehnder, Helio A. Costa

## Abstract

RNA sequencing is emerging as a powerful technique to detect a diverse array of fusions in human neoplasia, but few clinically validated assays have been described to date. We designed and validated a hybrid-capture RNAseq assay for FFPE tissue (Fusion-STAMP). It fully targets the transcript isoforms of 43 genes selected for their known impact as actionable targets of existing and emerging anti-cancer therapies (especially in lung adenocarcinomas), prognostic features, and/or utility as diagnostic cancer biomarkers (especially in sarcomas). 57 fusion results across 34 samples were evaluated. Fusion-STAMP demonstrated high overall accuracy with 98% sensitivity and 94% specificity for fusion detection. There was high intra- and inter-run reproducibility. Detection was sensitive to approximately 10% tumor, though this is expected to be impacted by fusion transcript expression levels, hybrid capture efficiency, and RNA quality. Challenges of clinically validating RNA sequencing for fusion detection include a low average RNA quality in FFPE specimens, and variable RNA total content and expression profile per cell. These challenges contribute to highly variable on-target rates, total read pairs, and total mapped read pairs. False positive results may be caused by intergenic splicing, barcode hopping / index hopping, or misalignment. Despite this, Fusion-STAMP demonstrates high overall performance metrics for qualitative fusion detection and is expected to provide clinical utility in identifying actionable fusions.

## Introduction

In human neoplasia, numerous clinically relevant translocations have been described, and more continue to be identified. Many are specific to one or several diagnoses, especially among soft tissue neoplasms. In conjunction with clinical history and histomorphologic/immunohistochemical findings, the detection of one of these translocations is a valuable diagnostic adjunct^1^. For example, in the setting of a small round blue cell tumor, translocation testing can help distinguish among differential diagnoses that include Ewing sarcoma, Ewing-like sarcomas, desmoplastic small round cell tumor, alveolar rhabdomyosarcoma, and synovial sarcoma, all of which are associated with distinct translocations or sets of translocations.

Other translocations may guide therapeutic decision making to optimally utilize targeted therapies, particularly in the setting of non-small cell lung carcinoma (NSCLC)^2^. For example, ALK, ROS1, and RET rearrangements are standard-of-care biomarkers predictive of a response to an FDA-approved medication in the setting of NSCLC. In addition, evidence is accumulating for clinical actionability of many other structural rearrangements in NSCLC and other tumors^3–5^.

Numerous techniques have been employed to detect fusions^3^. Traditional methods that do not employ next generation sequencing (NGS) include karyotyping, reverse transcriptase polymerase chain reaction (RT-PCR), and fluorescent in situ hybridization (FISH). Each of these methods has specific strengths and limitations. Karyotyping relies on growing cells in culture, can only detect large-scale alterations, demands significant interpretation time and can suffer from long turnaround time. RT-PCR is a sensitive and specific technique to test for well-characterized fusions with stereotyped breakpoints, but suffers from a limited ability to multiplex, or to detect novel rearrangements. FISH is considered the current gold standard for detecting fusions; though it greatly improves resolution compared to karyotyping, it still suffers from reduced sensitivity compared to NGS-based methods^6^, especially for small intrachromosomal events (“cryptic rearrangements”). Furthermore, FISH is unable to determine more granular details pertaining to fusions, including the fusion breakpoints, involved exons, and whether the fusion is in-frame or not. There is emerging evidence that these parameters may be clinically relevant. For example, in one reported cohort of patients with NSCLC positive by FISH testing for an EML4-ALK rearrangement and treated with ALK inhibitors, upon DNA and RNA NGS sequencing, patients with a predicted non-productive or no NGS-detectable EML4-ALK fusion demonstrated significantly worse mean survival compared to those with a predicted productive rearrangement^7^.

In more recent years, NGS-based fusion detection techniques have been developed. These include genomic DNA sequencing with target enrichment for regions in which breakpoints occur (such as selected “hotspot” introns)^8,9^, whole-transcriptome RNA sequencing utilizing poly(A) capture^10^, and targeted RNA sequencing employing hybridization-based capture^11,12^ or anchored multiplex PCR^13^. Broadly speaking, NGS-based techniques offer the advantage of greater breadth, depth, and resolution compared to traditional methods, with a tradeoff of increased cost.

Fresh tissue offers the best biospecimen quality characteristics for most molecular assays, but suffers from a lack of convenience, availability, and portability. In both clinical and research settings, formalin-fixed paraffin-embedded (FFPE) tissue has key advantages. These include being generated routinely in the clinical workflow and being a stable source of DNA and/or RNA for years after the tissue is acquired from the patient. In clinical practice, the need for fusion detection may not become apparent until after specimens are fixed and sections are examined under the microscope by a pathologist; also, the clinical need for fusion detection may change over time due to changes in the patient’s disease status, or evolution of knowledge in the field. However, FFPE presents significant biospecimen quality challenges to molecular assays due to chemical modifications including cross-linking which occur to DNA and RNA during fixation^14,15^. Cross-linking results in fragmentation, which limits the quantity of intact nucleic acids available for testing, and the obtainable length of NGS sequencing reads.

Each NGS-based fusion detection technique has advantages and limitations. Targeted DNA panels commonly used in cancer profiling can conveniently incorporate fusion detection by covering “hotspot” breakpoint regions and detecting fusion “spanning” or fusion “straddling” reads^8,9^. However, these panels can only capture a fraction of possible breakpoints, limited by intron sizes and fusion breakpoint diversity. Furthermore, targeted DNA panels on FFPE specimens have difficulties with repetitive or low complexity regions due to short read lengths; unfortunately, such regions often mediate genomic rearrangements^16^. On the other hand, RNA-based methods cannot detect rearrangements that do not lead to a fusion transcript, such as those that upregulate a gene’s expression by juxtaposing an enhancer element (eg, rearrangements involving IGH in some types of lymphoma), and may also miss lowly-expressed fusion transcripts. However, RNA-based NGS techniques can efficiently detect a diverse range of fusion breakpoints. Whole-transcriptome RNA sequencing using poly(A) capture on FFPE specimens for fusion detection has recently been reported^10^. This approach offers a wide breadth of sequencing and correspondingly a high discovery potential for novel fusions, which may be especially valuable in a research setting. However, due to RNA fragmentation, sensitivity decreases with the distance of the breakpoint from the poly(A) tail (ie breakpoints that are more 5’ in the fusion transcript suffer from reduced sensitivity)^10^. In addition, increased breadth of sequencing results in detection of more fusions of uncertain clinical significance. Some such fusions may be relevant but not yet understood, while others are likely to be “passenger fusions” which are not driving the cancer, but instead relate to copy number alterations or other structural alterations in cancers with genomic instability^17^. Whole-transcriptome sequencing also suffers from high cost and a prolonged turnaround time.

Currently, genes in which fusions are known to have clinical relevance comprise a small subset of the exome. A targeted panel enables optimization for cost-effective and sensitive detection of clinically relevant alterations. We have validated the Stanford Tumor Actionable Mutation Panel for Fusions (Fusion STAMP), a hybrid-capture based RNAseq assay (run on the Illumina MiSeq) that fully targets the transcript isoforms of 43 genes selected on the basis of their known impact as actionable targets of existing and emerging anti-cancer therapies, their prognostic features, and/or their utility as diagnostic cancer biomarkers. The targeted sequencing approach and integrated bioinformatics workflow is optimized for sequencing of FFPE tumor tissue specimens. In total, 34 unique samples (31 patient specimens, 1 purified RNA reference standard, and 2 RNA-FFPE reference standards [Horizon Discovery]) were tested in parallel by the Fusion STAMP method and compared to other reference methods to assess accuracy, yielding 57 fusion results. Reference methods included our in-house validated NGS panel for solid tumors (Stanford Actionable Mutation Panel; STAMP), validated fluorescence in situ hybridization (FISH) assays, and external reference testing performed by College of American Pathologists (CAP)-accredited laboratories. Analytical specificity was assessed using six non-neoplastic FFPE samples. Analytical sensitivity was assessed through serial dilution of an EWSR1 fusion cell line and by multiple analyses of the Seraseq Fusion RNA Mix v3 (SeraCare 0710-0431) which includes certified quantification of transcript levels by digital PCR. Intra-run, inter-run, and inter-instrument reproducibility was assessed. Here we describe the validation and anticipated clinical utility of Fusion STAMP.

## Materials and Methods

### Specimens and nucleic acid extraction

The patient tissue specimens described in this study were obtained from FFPE tissue blocks from Stanford Health Care. An anatomical pathologist reviewed, diagnosed, and estimated tumor purity from hematoxylin and eosin (H&E) slides of each specimen. A reference purified RNA sample (Seraseq Fusion RNA Mix v3; SeraCare, Cat. No. 0710-0431, Milford, MA, USA) harboring 14 known fusions targeted by our Fusion-STAMP panel was used as a positive control for our analyses. A reference FFPE sample with five fusion transcripts (EML4-ALK, CCDC6-RET, SLC34A2-ROS1, TPM3-NTRK1 and ETV6-NTRK3) confirmed to be present by endpoint RT-PCR (5-Fusion Multiplex (Positive Control) FFPE RNA Reference Standard; Horizon Discovery, Cat. No. HD796, Cambridge, UK) and a reference FFPE sample with the same five fusion transcripts confirmed to be absent by endpoint RT-PCR (5-Fusion Multiplex (Negative Control) FFPE RNA Reference Standard; Horizon Discovery, Cat. No. HD783, Cambridge, UK) were also tested. For dilution studies, a cell line containing an EWSR1 fusion (RD-ES (ATCC® HTB-166™); American Type Culture Collection (ATCC), Manassas, VA, USA) and a B-lymphocyte cell line from a patient with cystic fibrosis (GM07469; Coriell Institute for Medical Research, Camden, NJ, USA) were used. Total RNA from patient and control samples were extracted using a Qiagen RNeasy FFPE Kit (Qiagen Inc., Cat. No. 73504, Germantown, MD, USA), respectively. Additional specimen details can be found in **Table 2**.

**Table 1:**
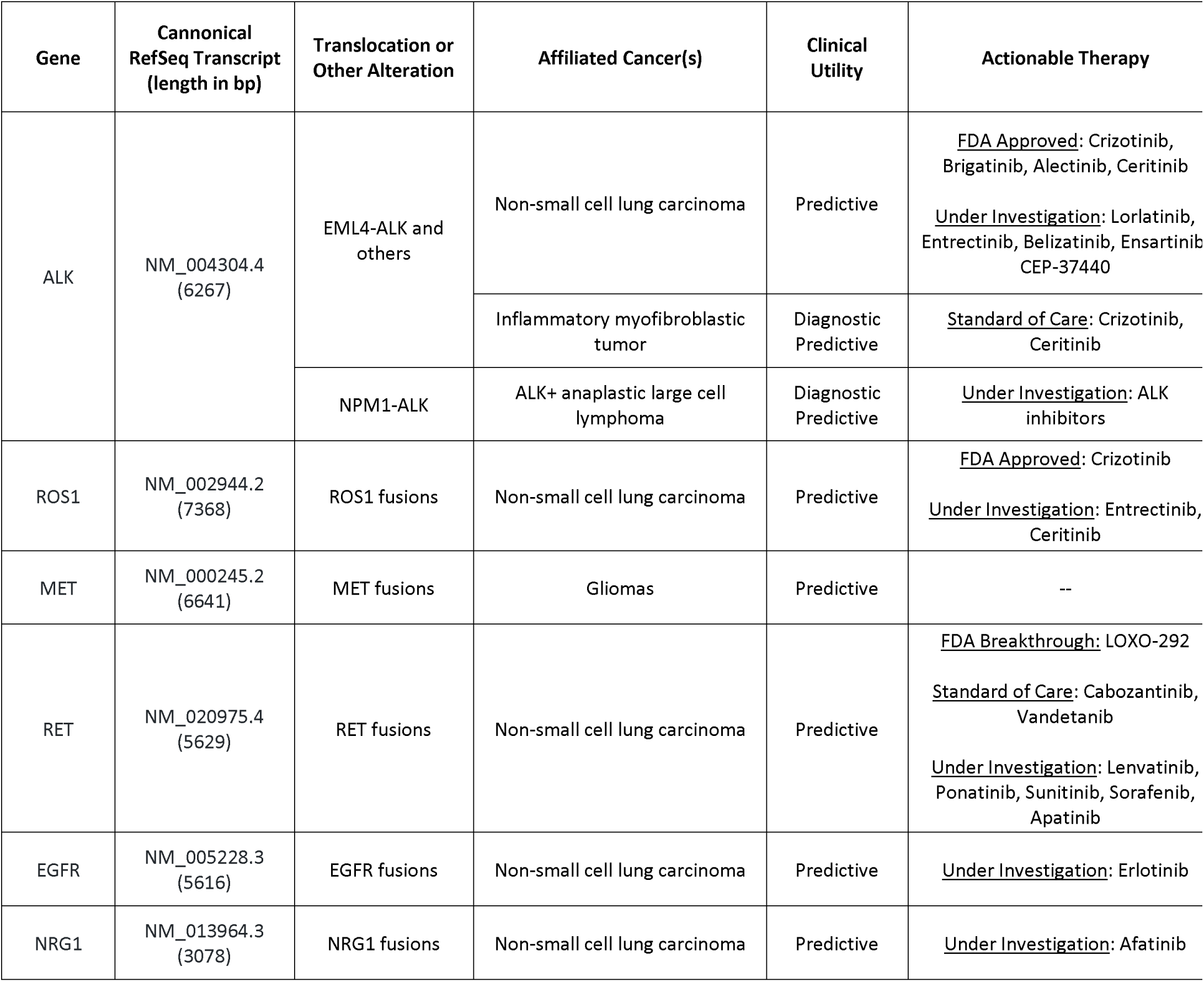
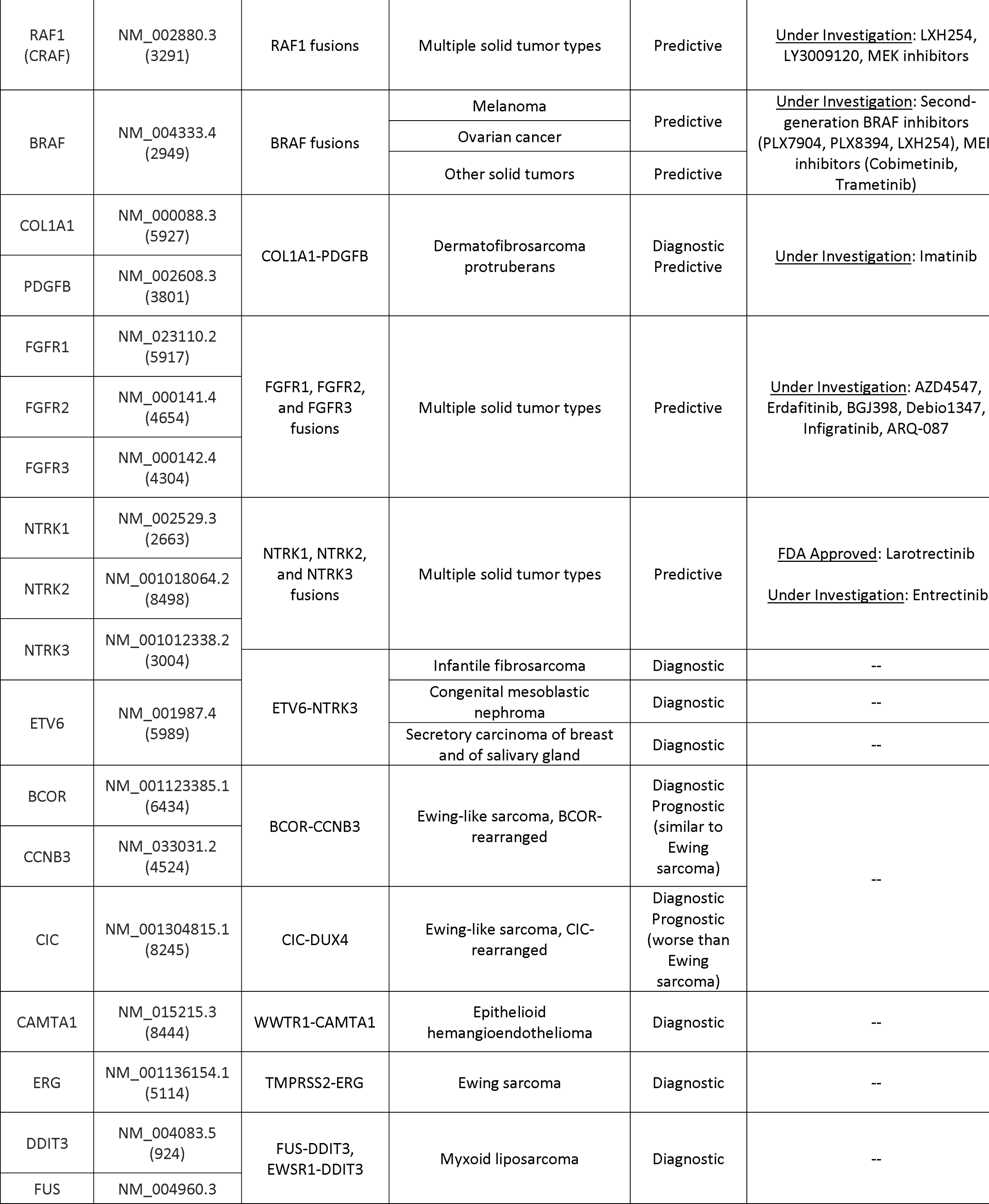
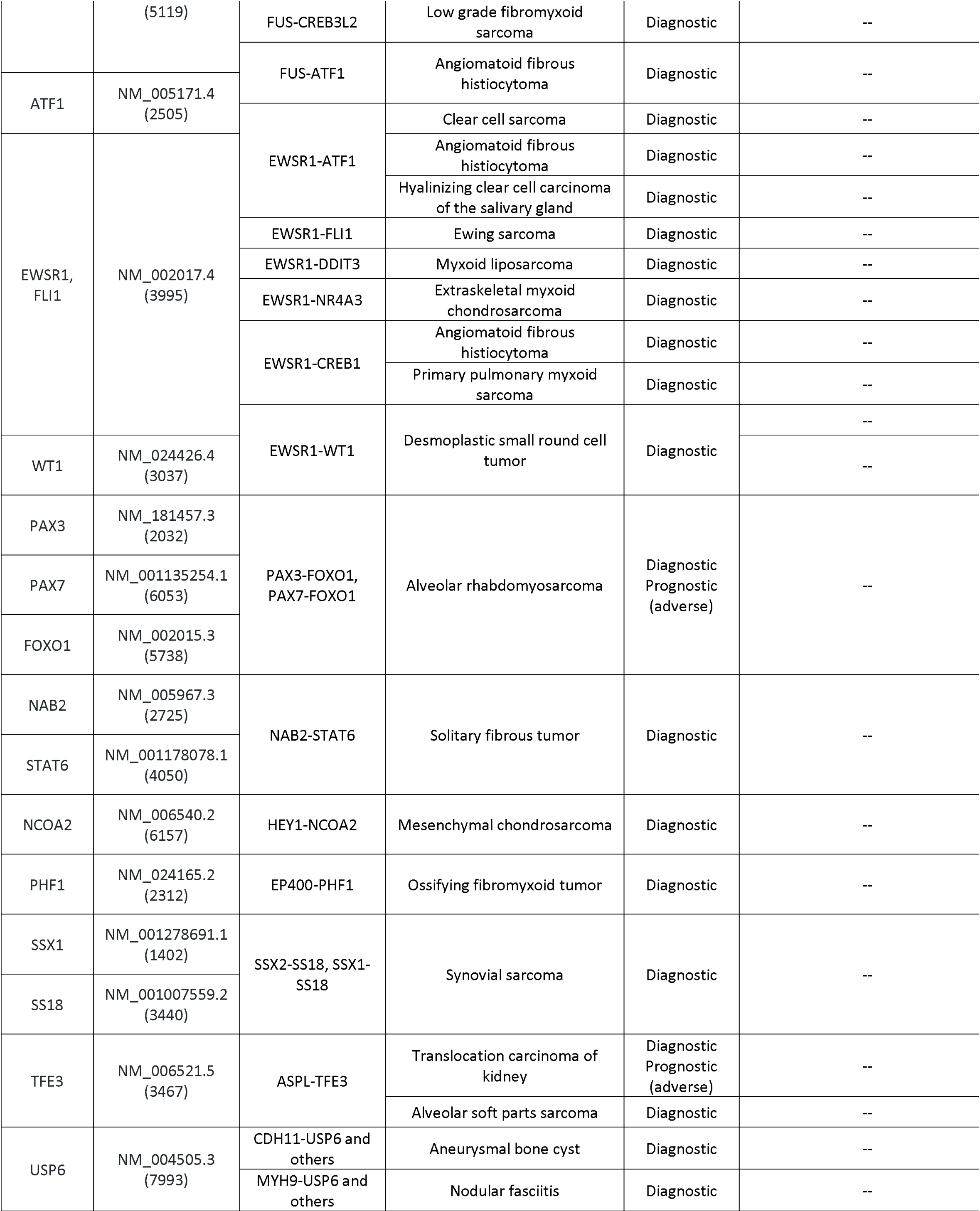
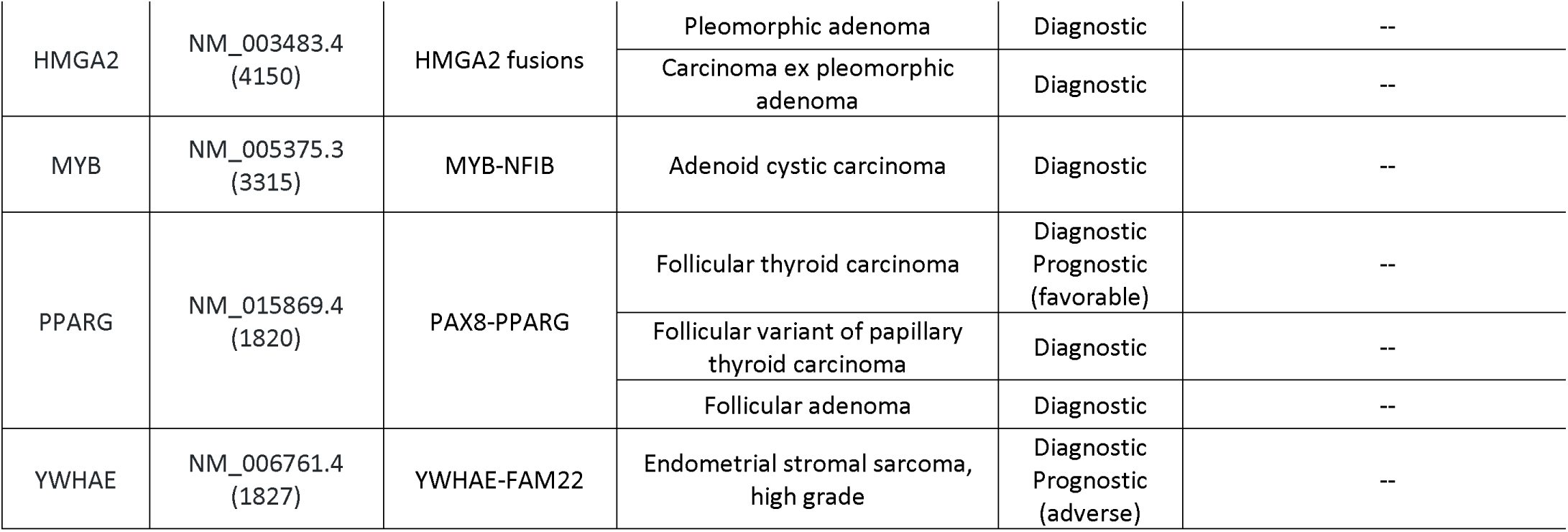
Fusion-STAMP gene panel with affiliated cancer(s) and clinical utility.

**Table 2:**
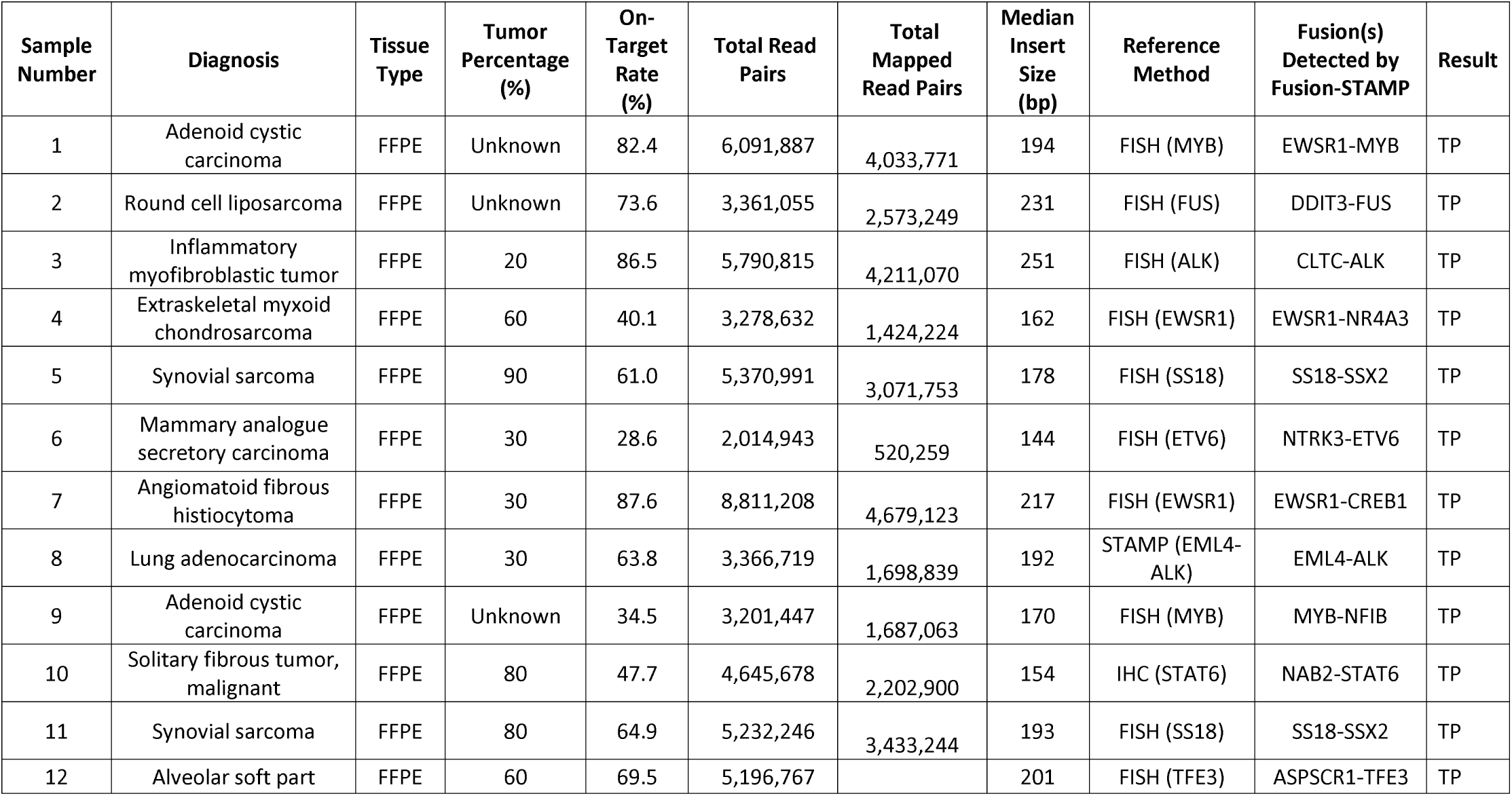
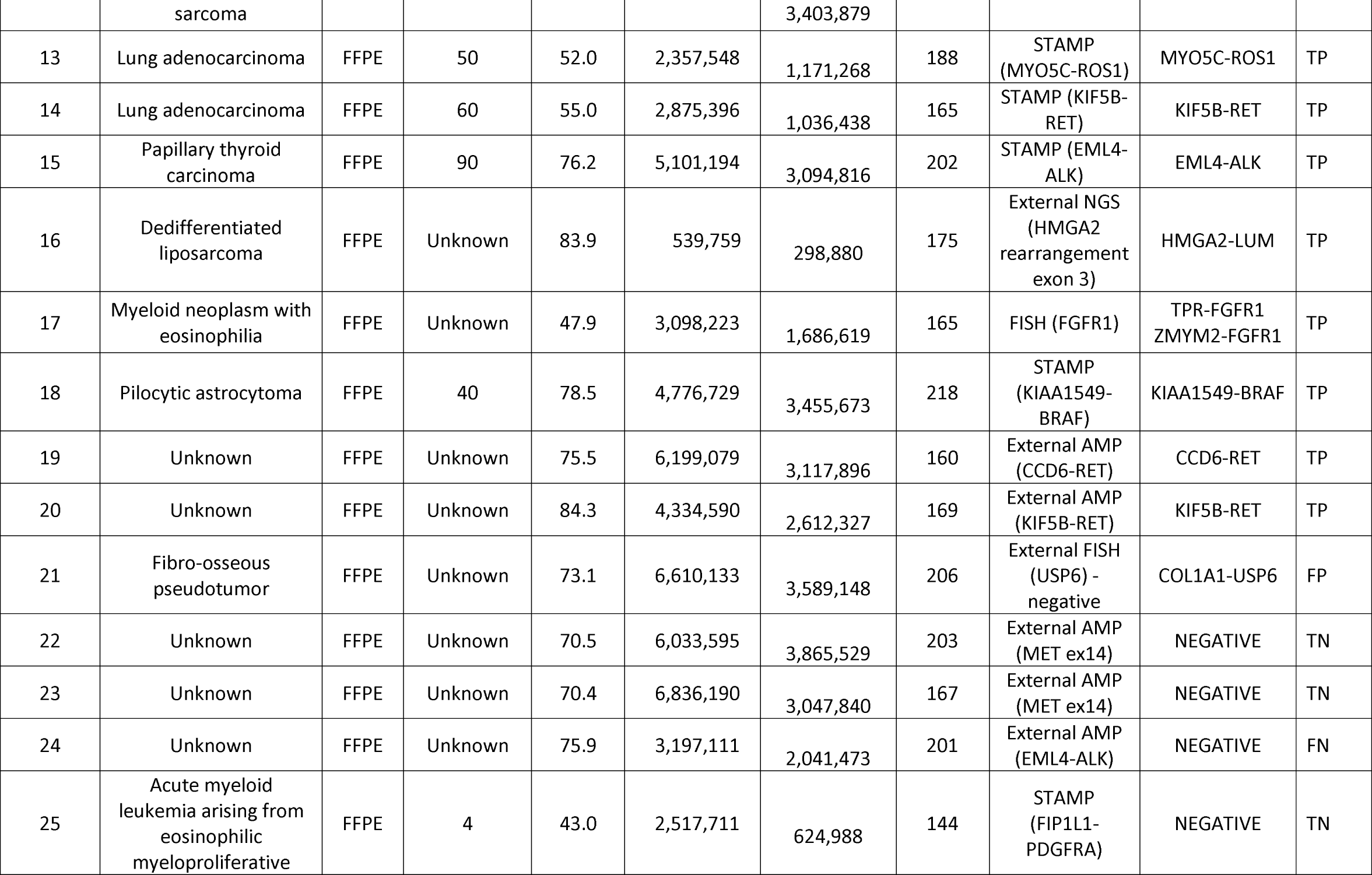
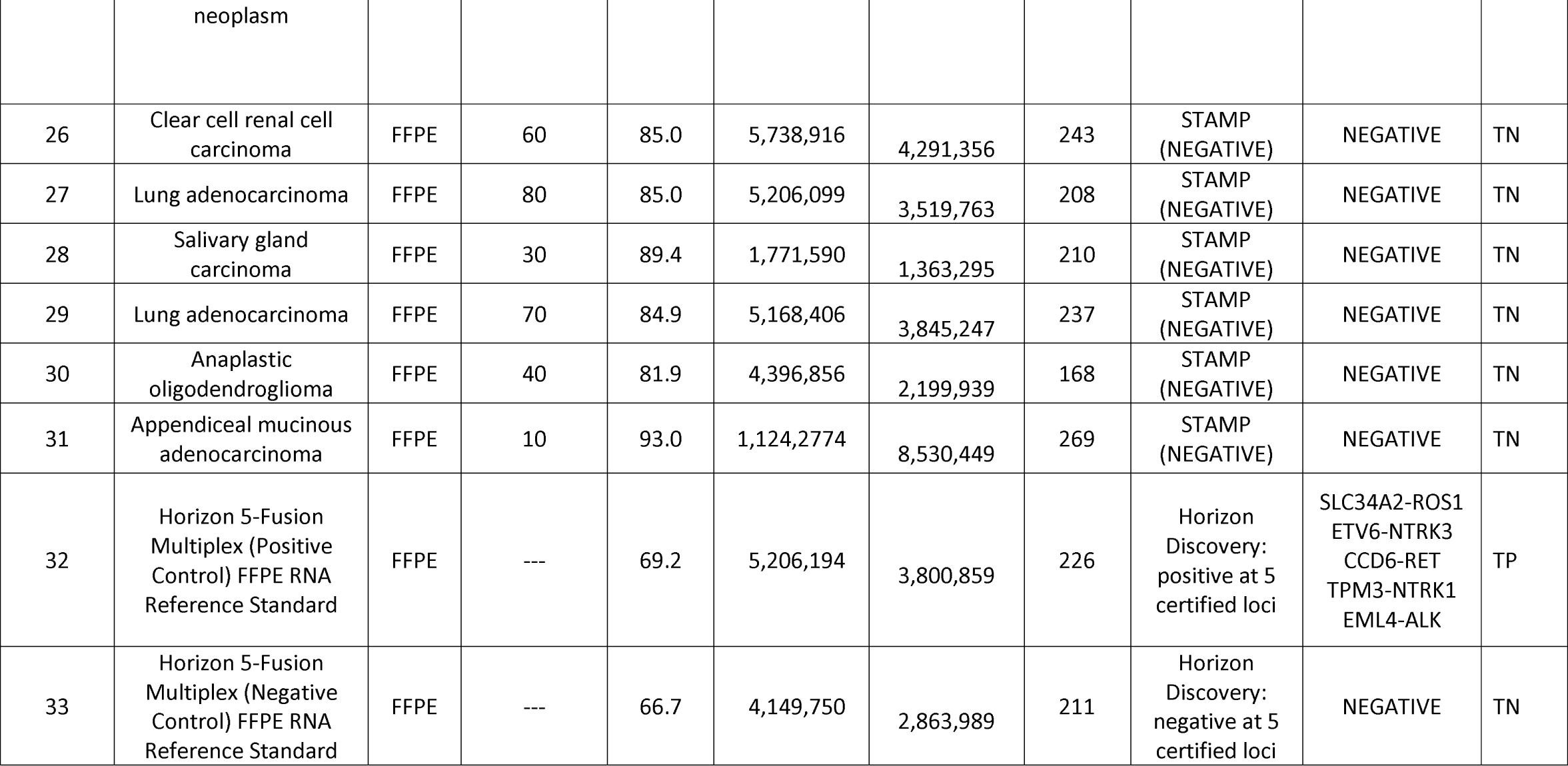
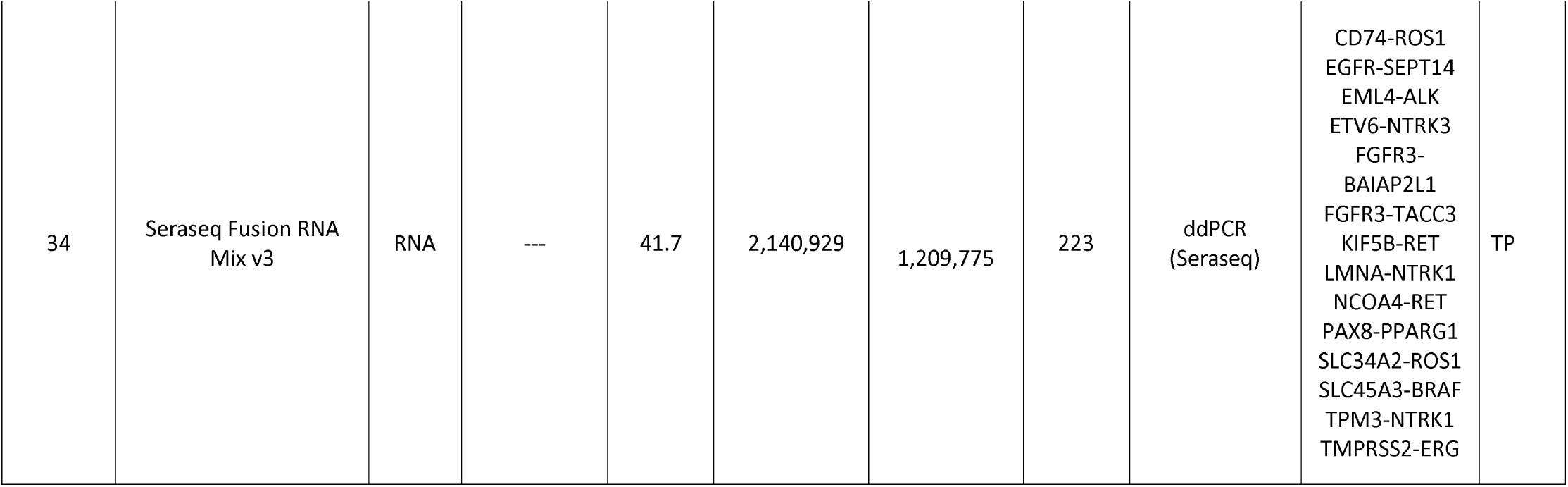
Sample pathological and sequencing metrics from Fusion-STAMP validation data. Abbreviations: FISH, fluorescent in situ hybridization; FFPE, formalin fixed paraffin embedded; STAMP, Stanford Actionable Mutation Panel; IHC, immunohistochemistry; AMP, anchored multiplex PCR; NGS, next generation sequencing; ddPCR, digital droplet polymerase chain reaction.

### Fusion-STAMP sequencing sample preparation, sequencing, and fusion detection

Total RNA (200ng input) from each specimen underwent cDNA synthesis and construction of sequencing libraries using a KAPA Stranded RNA-Seq Library Preparation Kit (Roche Sequencing, Cat. No. 07277261001, Pleasanton, CA, USA). Five to six samples at a time were then multiplexed and underwent enrichment for a 43-gene targeted RNA fusion panel (Table 1) using Roche SeqCap RNA Choice target enrichment probes spanning the entirety of the gene transcripts of interest (Roche Sequencing, Cat. No. 6953247001, Pleasanton, CA, USA). Sequencing was then performed on an Illumina MiSeq instrument producing 150bp paired end reads. In brief, sequencing reads were mapped to the human reference genome (GRCh38.p12) using the STAR-Fusion algorithm (v 1.1.0). STAR-Fusion uses the STAR aligner^18^ to map reads and identify candidate fusion transcripts, which are then processed by the STAR-Fusion algorithm to map junction reads and spanning reads to a reference annotation set and to produce a final fusion transcript list. STAR-Fusion is run with the following parameters: STAR-Fusion_v1.1.0/STAR-Fusion --left_fq <R1.fastq.gz> --right_fq <R2.fastq.gz> --genome_lib_dir <genome reference directory> --FusionInspector validate --annotate examine_coding_effect --extract_fusion_reads. Called variants were annotated for a series of functional predictions using publicly available database annotations via internal perl scripts.

### Fluorescent In Situ hybridization (FISH)

FISH analysis was performed on interphase nuclei or metaphase chromosomes with the corresponding break-apart FISH probe as previously described^19^.

### Statistical analyses

All statistical analyses were performed in the R programming language.

## Results

### Overview of Gene Panel Targets

Forty-three genes were targeted by Fusion STAMP based on literature review of clinical utility. Of these, 15 are protein kinases involved in fusions with established or emerging evidence for clinical actionability with targeted therapies^3^; 31 are involved in fusions with diagnostic utility; and 9 are involved in fusions with prognostic utility (Table 1). Some targeted genes are involved in fusions with multiple domains of clinical utility. For example, PAX3-FOXO1 and PAX7-FOXO1 are diagnostic for alveolar rhabdomyosarcoma, and also portend a worse prognosis (especially PAX3-FOXO1) compared to embryonal rhabdomyosarcoma and fusion-negative alveolar rhabdomyosarcoma^20^. Some fusions have differing clinical significance depending on the tumor type and the translocation partner; for example, NTRK3 fusions occur across many solid tumor types and may predict response to targeted therapies including larotrectinib and entrectinib^3^, but ETV6-NTRK3 fusions are diagnostic markers for infantile fibrosarcoma, congenital mesoblastic nephroma, and secretory carcinoma of the breast and salivary gland. Overall the selected genes provide clinical utility across multiple scenarios. These include NSCLC, particularly those negative for a typical MAPK pathway driver mutation; sarcomas, especially small round blue cell tumors, or others that are difficult to classify; and select head and neck entities, including certain thyroid and salivary gland tumors.

### Overview of Experimental and Computational Workflow

The Fusion STAMP workflow includes isolation of total RNA molecules, followed by efficient preparation of sequencing libraries and a target enrichment approach to capture mRNA transcript regions of interest for sequencing. The enrichment is done using custom designed libraries of capture oligonucleotides that target a specific set of expressed genomic regions. This panel fully targets the major canonical transcript isoforms of the 43 genes described above. The bioinformatic pipeline includes sequencing quality control, paired-end mapping to the human transcriptome, and detection of gene fusion events using the STAR-Fusion algorithm. In addition, quality control metrics and plots are generated from the aligned BAM files. A molecular genetic pathology fellow or clinical molecular genetics fellow reviews all fusion variant calls.

### Sequencing Metrics and Clinical Reporting Thresholds

Sequencing metrics across the 34 tested samples (Table 2) demonstrate significant variability in on-target rate (range: 28.6 - 93.0%), total read pairs (range: 539,759 – 11,242,774), mapped read pairs (range: 298,880 – 8,530,449), and insert size (range: 144 – 269 bp) despite uniform input RNA mass (200 ng) for all specimens.

True positive fusions had variable read support in the validation cohort, sometimes <20 junction reads (i.e. a read that aligns as a split read at the site of the putative fusion junction). Also, in many samples, low numbers of junction reads (generally <20) were identified for fusions which did not fit in the clinical or biological context, and had not been previously reported in the literature. These were often adjacent or nearby in the genome (possibly representing intergenic splicing^21^). Based on the levels of junction read support for true positive fusions and this presumed noise in the validation data, clinical reporting thresholds were set to optimize performance metrics in the validation cohort. A “whitelist” was created of fusions which have previously been reported in the literature, and a lower reporting threshold was set for these fusions. A higher threshold was set for fusions with identical breakpoints to one or more other multiplexed samples, due to the phenomenon of barcode hopping^22^. For clinical testing, fusions with supporting junction reads below the reporting threshold that are suspected of being diagnostically or clinically significant may be confirmed by RT-PCR and Sanger sequencing, or by another corroborating result (such as FISH).

### Analytical Sensitivity / Limit of Detection, and Analytical Specificity

Analytical sensitivity was assessed with six replicates of the Seraseq Fusion RNA Mix v3, which contains 14 fusion variants whose presence is confirmed and quantitated by digital PCR. Supporting junction reads for 13 of the 14 fusions were detected in all replicates. There was not a clear proportional relationship between the ddPCR copy number as reported by SeraCare (which is based on the number of supportive reads with unique start sites), and the number of supporting junction reads detected on Fusion STAMP. One fusion (TMPRSS2-ERG) was not detected in one replicate (1/6; 17%), and junction read support for this fusion was low in the other replicates. Since this fusion appeared to be near the limit of detection of the Fusion STAMP assay, detection of this fusion was dropped from QC requirements for clinical testing.

Limit of detection was further assessed with a cell line dilution study. Single-replicate serial dilutions were performed using a cell line with an EWSR1 fusion, and a cystic fibrosis cell line (Figure 2). Junction read support of ≥20 reads was demonstrated down to a dilution of 6.25%. Of note, 3 junction reads were detected in the 100% cystic fibrosis cell line sample. This is below the established reporting threshold and may suggest barcode hopping or trace contamination. Overall, based on this data, the sensitivity of Fusion STAMP is cited as approximately 10% tumor.

**Figure 1:**
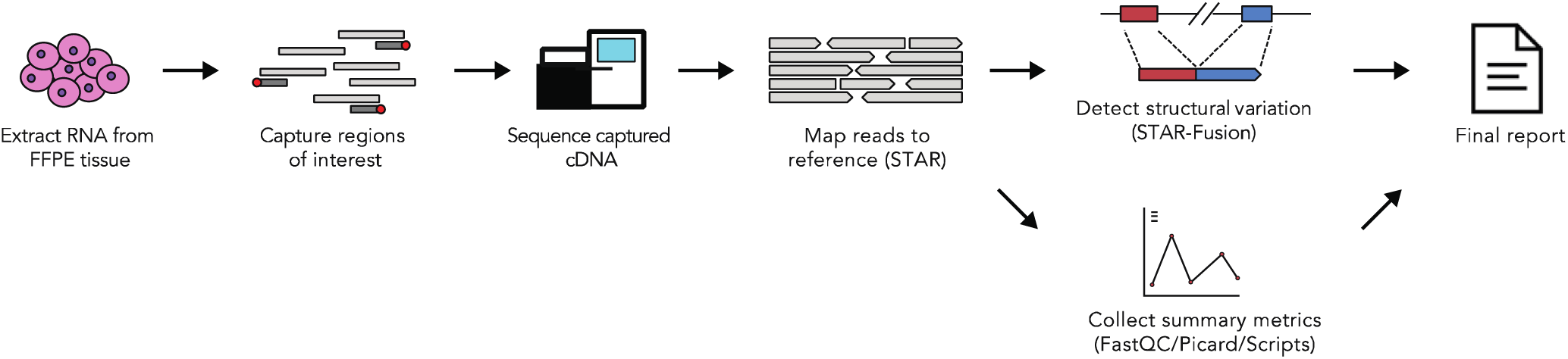
Fusion-STAMP experimental and computational workflow.

**Figure 2:**
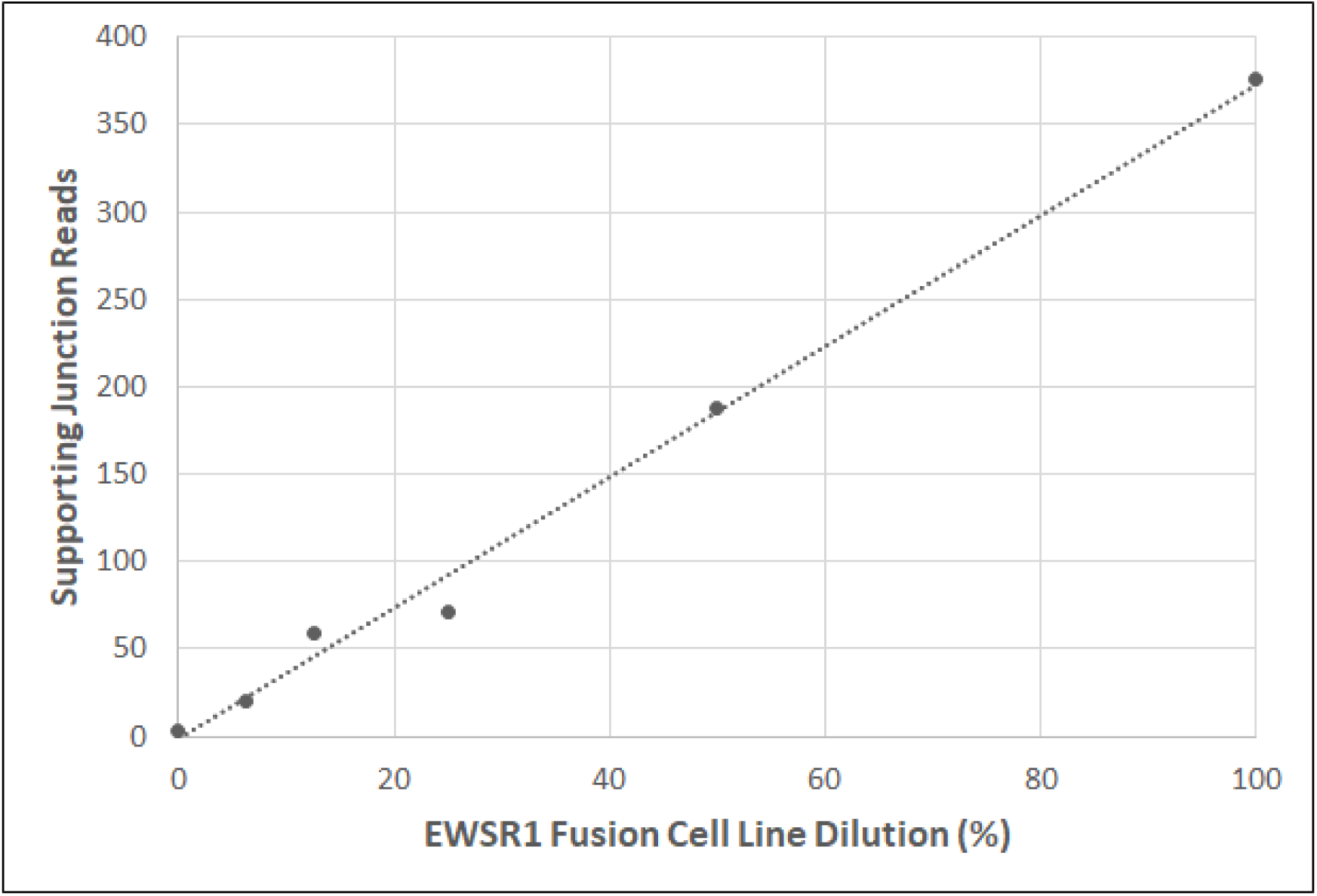
Fusion-STAMP limit of detection study. Single-replicate serial dilutions were performed using a cell line with an EWSR1 fusion, and a cystic fibrosis cell line.

Analytical specificity was assessed by testing 6 non-neoplastic tissue FFPE specimens. No fusions were detected above the reporting threshold in these samples.

### Reproducibility (Precision)

Intra-run and inter-run reproducibility were assessed in three replicates each of the Seraseq Fusion RNA Mix v3 and two clinical samples, one with an EML4-ALK translocation, and the other with a MYO5C-ROS1 translocation. All fusions in the Seraseq control (excluding the TMPRSS2-ERG fusion), and the EML4-ALK and MYO5C-ROS1 fusions, were detected across all replicates.

### Accuracy

Fusion STAMP showed excellent accuracy (Table 3). One case was negative by FISH for USP6 but Fusion STAMP was positive for a COL1A1-USP6 fusion. Given the tumor context (a fibro-osseous pseudotumor of the digit), this likely represents a false negative result by FISH testing. Fusion-STAMP is expected to show greater analytical sensitivity than FISH.

**Table 3:** Summary of accuracy testing metrics. Each fusion call, and each negative sample, was counted as one call. Abbreviations: TP, true positive; FP, false positive; FN, false negative; TN, true negative; Sn, sensitivity; Sp, specificity; PPV, positive predictive value; NPV, negative predictive value.

|  | Fusion-STAMP Positive | Fusion-STAMP Negative | Metric |
| --- | --- | --- | --- |
| Reference Method Positive | 40 (TP) | 1 (FN) | Sn: 98% |
| Reference Method Negative | 1 (FP) | 15 (TN) | Sp: 94% |
| Metric | PPV: 98% | NPV: 94% | Total:<br>57 calls |

One case was positive by outside testing for an EML4-ALK rearrangement, but this was not detected by Fusion STAMP. In this case, the outside lab performed micro-dissection for tumor enrichment. This was not an option with the material received for Fusion STAMP; this may account for this false negative result.

Of the 43 genes in this panel, 27 are involved in at least one fusion in the validation data set. Of the remaining 16 genes, many are rarely involved in fusions, making it a challenge to obtain reference material. To demonstrate that the selector was successful in capturing these transcripts when expressed, we examined the coverage data in appropriate tissue types among our validation samples. Demonstrable capture was identified for all transcripts on the selector in at least one sample.

## Discussion

Though RNAseq on FFPE promises multiple advantages over DNA sequencing, it also comes with numerous challenges. This includes a low average RNA quality in FFPE specimens, and variable RNA total content and expression profile per cell. The downstream effects of these issues can be seen in highly variable on-target rates, total read pairs, and total mapped read pairs in the Fusion STAMP validation cohort. Tumor percentage estimates, while still important, are less directly related to the fraction of RNA read pairs that originate from the tumor than they would be for DNA. It is conceivable that a lowly expressed fusion could be missed despite relatively high tumor percentage, especially in a poor-quality specimen. Also, given the multiplex design, even though the hybrid capture input RNA mass per sample is constant, variable expression profiles between samples can result in disproportionate sequencing of some samples with greater RNA content aligning to the selector at the expense of other samples.

The sensitivity of Fusion STAMP is estimated to be around 10% tumor based on the EWSR1 cell line dilution study performed during validation; however, this sensitivity is expected to vary significantly by the hybrid capture efficiency of the involved genes, the fusion transcript expression level, and the specimen quality. One false negative was identified in the validation cohort and appears likely to relate to low tumor percent due to lack of enrichment, and poor RNA quality. However, Fusion STAMP demonstrated high sensitivity for fusion detection overall.

False positive results may be caused by intergenic splicing^21^, barcode hopping / index hopping^22^, or misalignment. These findings are expected to vary depending on the expression profile of the sample, and therefore will likely vary by the site of origin of the tissue. The full range of human tissue types is near-impossible to comprehensively assess during validation. Several tissue types were tested during validation including lung, gastrointestinal tract, and soft tissue, and no false positives were detected above the reporting thresholds. As clinical testing continues and more tissue types are sequenced, recurrent artifacts will be prospectively tracked, identified, and/or filtered.

Multiple RNA NGS sequencing quality control strategies and metrics have been described in the literature. These include spike-in control transcripts and corresponding probes to assess efficiency of hybrid capture and indirectly assess RNA quality^11^; probes to RNA from housekeeping genes to assess RNA quality^11^; a minimum total mapped read count^10,11,13^; a minimum on-target rate^11^; a minimum mapped exon-exon junction read count^10^; a percent of mapped reads that map to coding regions^10^; and qPCR-based assessment of RNA quality^13^. The utility of these metrics needs to be weighed against the theoretical possibility of detecting a highly expressed fusion despite poor quality, or of missing a lowly expressed fusion despite high quality. This makes it challenging to have an accurate assessment of the risk of a false positive or negative result in any individual case. For this Fusion STAMP validation cohort, despite employing only run-level QC criteria and sample-specific total mapped reads QC cutoffs, after optimizing reporting cutoffs to exclude noise and include real events as confirmed by ancillary testing, the cohort demonstrates a high sensitivity, specificity, precision and accuracy for qualitative fusion detection.

## Conclusions

Fusion STAMP is a hybrid-capture based RNAseq assay (run on the Illumina MiSeq) that fully targets the transcript isoforms of 43 genes selected on the basis of their known impact as actionable targets of existing and emerging anti-cancer therapies, their prognostic features, and/or their utility as diagnostic cancer biomarkers. Despite challenges related to sequencing RNA from FFPE tissue, after optimizing cutoffs to exclude noise and include real events, this validation cohort demonstrates a high sensitivity, specificity, precision and accuracy for fusion detection. This assay is expected to provide clinical utility in the setting of NSCLC, particularly those negative for any known driver mutation; sarcomas, especially small round blue cell tumors, or others that are difficult to classify; and select head and neck entities, including certain thyroid and salivary gland tumors.

## Acknowledgements

All authors contributed to the study design. C.J., S.S., H.Z. and H.A.C. performed the benchwork experiments. E.F. and H.A.C. performed the computational analyses. All authors contributed to the final draft of the manuscript. H.A.C. is the guarantor of this work, and, as such, had full access to all of the data in the study and takes responsibility for the integrity of the data and the accuracy of the data analysis.

